# Critical importance of DNA binding for CSL protein functions in fission yeast

**DOI:** 10.1101/2023.08.22.554308

**Authors:** Anna Marešová, Martina Oravcová, Miluše Hradilová, Viacheslav Zemlianski, Robert Häsler, Martin Převorovský

## Abstract

CSL (CBF1/RBP-Jκ/Suppressor of Hairless/LAG-1) proteins are conserved transcription factors found in animals and fungi. In fission yeast, they regulate various cellular processes, including cell cycle progression, lipid metabolism, and cell adhesion. CSL proteins bind to DNA through their N-terminal Rel-like domain and central beta-trefoil domain. Here, we investigated the importance of DNA binding for CSL functions in the fission yeast *Schizosaccharomyces pombe*. We created CSL mutants with disrupted DNA binding and found that the vast majority of CSL functions depend on intact DNA binding. Specifically, DNA binding is crucial for the regulation of cell adhesion, lipid metabolism, cell cycle progression, long non-coding RNA expression, and genome integrity maintenance. Interestingly, perturbed lipid metabolism leads to chromatin structure changes, potentially linking lipid metabolism to the diverse CSL-associated phenotypes. Our study highlights the critical role of DNA binding for CSL protein functions in fission yeast.

**SUMMARY STATEMENT:** CSL transcription factors regulate a diverse set of processes, but the mechanisms are not always clear. We show that *S. pombe* CSL proteins need the ability to bind DNA for most of their roles.

## INTRODUCTION

CSL (CBF1/RBP-Jκ/Suppressor of Hairless/LAG-1) proteins, a structurally unique and conserved family of transcription factors, are found in animals and fungi (with the exception of most Ascomycetes) (Hall and Kovall, 2019; Převorovský et al., 2007). Their regulatory roles in animal development have been studied extensively (for a review, see e.g. (Borggrefe and Oswald, 2009)), while the fungal CSL proteins have only been characterized in the fission yeast *Schizosaccharomyces pombe*, where they regulate cell cycle progression, lipid metabolism, and cell adhesion (Kwon et al., 2012; Prevorovský et al., 2009; Převorovský et al., 2015; Vachon et al., 2013). The mechanism of CSL protein binding to DNA has been characterized in detail, and is mediated by the N-terminal Rel-like domain (NTD) together with the central beta-trefoil domain (BTD) (Kovall and Hendrickson, 2004). Notably, the DNA binding mode seems to be conserved throughout evolution, as BTD mutations disrupting DNA binding in the murine RBP-Jκ protein (substitution R218H; (Chung et al., 1994)) also abolish binding to DNA of fission yeast Cbf11 (substitution R318H) and Cbf12 (substitution R644H) CSL paralogs (Oravcová et al., 2013).

CSL proteins were shown to participate in numerous processes in *S. pombe*, with Cbf11 often acting as an antagonist of Cbf12 (Prevorovský et al., 2009). For example, Cbf12 is required for cell-surface and cell-cell adhesion (non-sexual flocculation). Cells lacking Cbf12 lose their ability to adhere, while overexpression of Cbf12 leads to a hyper-adhesive phenotype. On the contrary, cells lacking Cbf11 show increased adhesion, and adhesion is abolished upon overexpression of Cbf11 (Kwon et al., 2012; Prevorovský et al., 2009). Later, it was shown that CSL proteins are part of an elaborate network of several transcription factors which regulate flocculation by activating or repressing genes encoding adhesive cell-surface proteins (flocculins *gsf2*, *pfl2*-*pfl9*; (Kwon et al., 2012; Linder and Gustafsson, 2008)) and cell wall-remodeling enzymes (Kwon et al., 2012). Cbf12 is upregulated in the stationary phase of growth (Prevorovský et al., 2009), when cell adhesion naturally increases, and Cbf12 can activate its own transcription in a positive feedback loop (Kwon et al., 2012).

Likewise, the transcriptional regulation of lipid metabolism genes in *S. pombe* is controlled by a network of at least three transcription factors. The metabolism of triglycerides is controlled by the transcription factors Cbf11 and Mga2, which share some target genes (Burr et al., 2016; Převorovský et al., 2015; Převorovský et al., 2016). On the other hand, the metabolism of sterols is controlled by Sre1 (mammalian SREBP-2 homolog), and there is crosstalk between Sre1 and Mga2 that coordinates the two branches of lipid metabolism (Burr et al., 2016; Burr et al., 2017). Intriguingly, *S. pombe* contains another SREBP paralog, Sre2, which does not seem to regulate lipid metabolism, but is involved in regulating flocculation (Kwon et al., 2012). This is reminiscent of the relationship between the Cbf11 and Cbf12 CSL paralogs (Prevorovský et al., 2009). The target genes of Cbf11 include the *cut6* acetyl-coenzyme A carboxylase (the rate-limiting enzyme of fatty acid synthesis), *ole1* acyl-coenzyme A desaturase, long-chain fatty acid-coenzyme A ligases *lcf1* and *lcf2*, and *bio2* biotin synthase and *vht1* biotin transporter (biotin is a cofactor for Cut6) (Převorovský et al., 2015).

Cells lacking *cbf11* or overexpressing *cbf12* also display a range of defects related to cell cycle progression (Prevorovský et al., 2009). Most notably, CSL manipulations lead to increased incidence of a particular form of catastrophic mitosis (“cut” phenotype; (Prevorovský et al., 2009; Yanagida, 1998)), a phenotype shared with several other lipid-related mutants (Zach and Převorovský, 2018). It was suggested that, when fatty acid production is inhibited, there is insufficient supply of phospholipids for nuclear envelope expansion during the anaphase of the closed mitosis in *S. pombe*, and this results in spindle collapse and mitotic failure (Makarova et al., 2016; Takemoto et al., 2016). Moreover, we recently identified additional factors contributing to mitotic defects in *Δcbf11* cells, such as perturbed structure of centromeric chromatin and aberrant cohesin occupancy (Vishwanatha et al., 2023). However, the molecular details of how dysregulated lipid metabolism leads to these various cell cycle defects remain elusive. Interestingly, lipid metabolism has a pronounced impact on chromatin structure and proper regulation of stress gene expression. For example, cells expressing the Cbf11DBM mutant protein with abolished DNA binding (the above-mentioned R318H substitution) show wide-spread changes in histone H3 lysine 9 (H3K9) modifications, including specific H3K9 hyperacetylation and derepression of oxidative stress response genes, which is mediated by the histone acetyltransferases Gcn5 and Mst1 (Princová et al., 2023).

Transcription regulators can control their target genes by binding DNA either directly, or indirectly as part of a protein complex. There are also examples of transcription factors with moonlighting roles, completely independent of their transcription factor functions (Lee et al., 2003; Wegrzyn et al., 2009). Furthermore, mutated mammalian CSL proteins unable to bind DNA have been shown to affect signaling and gene expression by sequestering transcriptional coregulators away from DNA (Gagliani et al., 2022; Kato et al., 1997). To better understand the functional relationships within the broad spectrum of seemingly disparate roles of CSL proteins in *S. pombe*, we set out to determine the requirements for DNA binding for each known CSL role. In this study we show that, rather unexpectedly, the vast majority of CSL functions in *S. pombe* cells depend on their intact DNA binding domain (BTD domain). We also hypothesize that changes in chromatin structure resulting from perturbed lipid metabolism could be the unifying theme behind the diversity of CSL-associated phenotypes.

## RESULTS

### Creation of CSL DNA binding mutant strains

To determine the importance of DNA binding activity for the various roles of fission yeast CSL proteins we employed previously described point mutations that disrupt DNA binding (DNA-binding mutation, DBM, (Oravcová et al., 2013)). We re-created the DBM mutations directly in the endogenous *cbf11* and *cbf12* loci, respectively, and added a C-terminal TAP-tag to facilitate protein detection and purification. To introduce as few confounding factors as possible, the whole construction was performed in a scarless manner (see Materials and Methods for details), leaving no selection markers and keeping the regulatory untranslated regions intact (Wells et al., 2012). As controls, we also created TAP-tagged alleles of non-mutated *cbf11* and *cbf12* in an analogous manner. The introduction of the TAP tag did not affect growth rate. Notably, the growth of the DBM mutant strains was comparable to growth of the respective deletion mutants (**Fig. S1**).

### DNA-binding activities of CSL proteins are required for their antagonism in cell adhesion

Cbf11 and Cbf12 antagonistically regulate adhesion to agar and cell flocculation (Prevorovský et al., 2009), with Cbf12 directly activating the expression of several adhesion-related genes, including the *cbf12* gene itself (Kwon et al., 2012). Notably, *cbf11DBM-TAP* cells displayed increased adhesion to agar comparable to the *Δcbf11* strain (**Fig. 1A**), while the adhesion of *cbf12DBM-TAP* cells was abolished as in the *Δcbf12* strain (**Fig. 1B, Fig. S2**). These results indicate that the DNA binding activity is required for both CSL proteins to exert their regulatory functions in cell adhesion.

**Figure 1:**
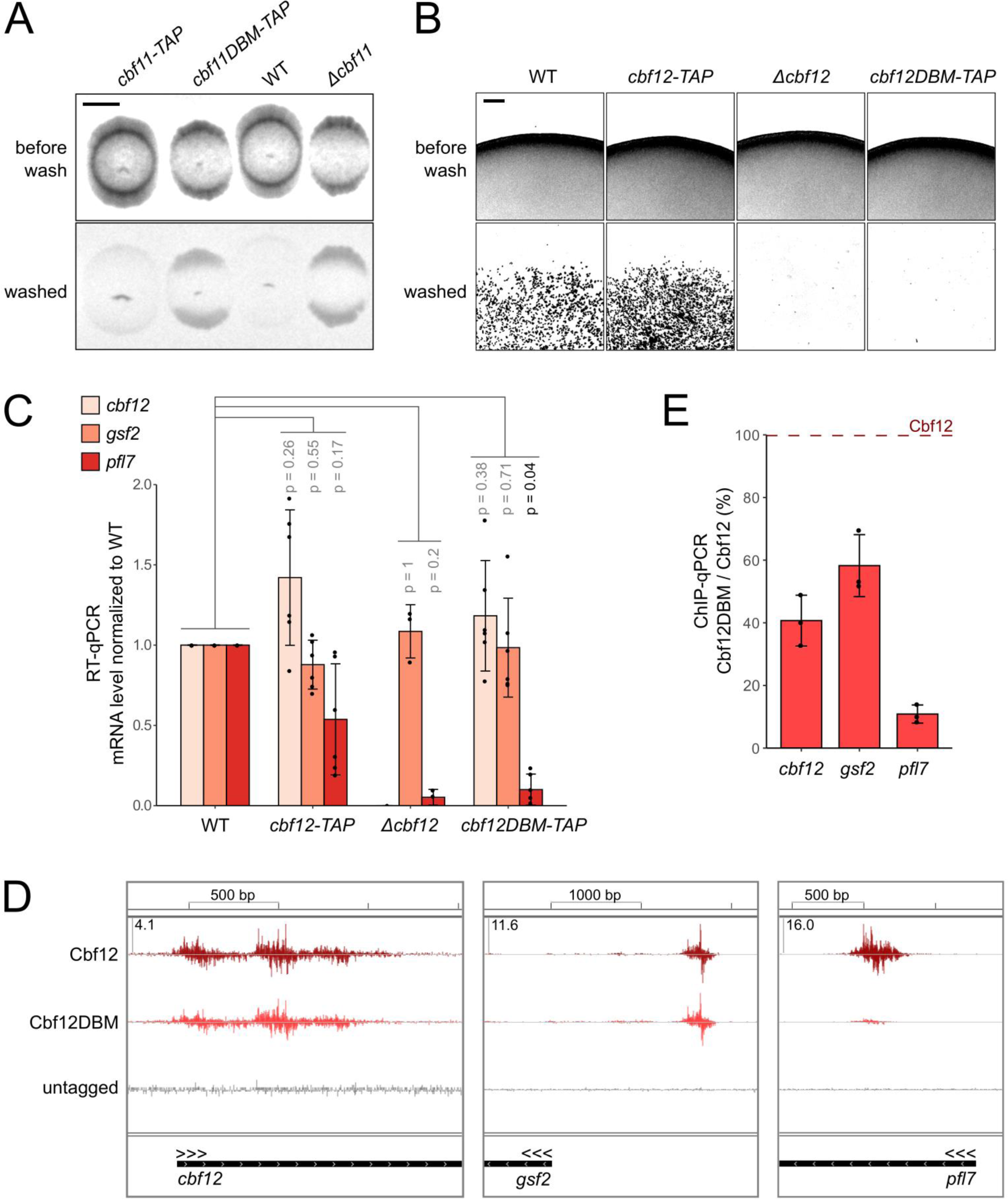
DNA-binding activities of CSL proteins are required for their antagonism in cell adhesion. **A)** Both *Δcbf11* and *cbf11DBM-TAP* cells display higher adhesion to agar than WT and *cbf11-TAP* cells. Cells were spotted on a plate, incubated at 32°C for 8 days, and then washed with a stream of water. Whole spots were imaged before and after washing. Representative images of 2 independent repeats are shown. Brightness and contrast were adjusted for clarity. Scale bar represents 5 mm. **B)** Cells lacking *cbf12* as well as *cbf12DBM-TAP* cells do not adhere to agar. Cells growing for 7 days on a plate were subjected to a washing test. A microscopic picture of the colony edge was taken before and after washing. Representative images of 3 independent experiments are shown. Brightness and contrast were adjusted for clarity (original image is shown in **Fig. S2**). Scale bar represents 100 μm. **C)** Expression of the indicated adhesion-related genes in WT, *cbf12-TAP*, *Δcbf12*, and *cbf12DBM-TAP* cells grown to stationary phase was analyzed by RT-qPCR. Mean ± SD values, as well as individual data points for ≥2 independent experiments are shown. Two-sided Mann-Whitney U test was used to determine statistical significance. Note that the change of *pfl7* mRNA level in *Δcbf12* is not statistically significant because only two measurements were done. **D)** *In vivo* binding of Cbf12 and Cbf12DBM TAP-tagged proteins to the promoters of indicated adhesion genes in the stationary phase was analyzed by ChIP-nexus. Mean strand-specific coverage profile of 3 independent experiments for Cbf12 and Cbf12DBM, and a strand-specific coverage profile for untagged WT cells (negative control) were visualized in the Integrated Genome Viewer (IGV; Broad Institute). Gene orientation is indicated by three arrowheads; all three sample tracks have the same Y-axis scaling, with the maximum Y-axis value for each locus given in the top-left corner. **E)** ChIP-qPCR shows the decrease in DNA binding of the Cbf12DBM-TAP protein compared to its unmutated counterpart at selected target loci in the stationary phase. Mean ± SD values, and individual data points of 3 independent replicates are shown.

To study the effect of DBM mutations in more detail, we first performed a genome-wide ChIP-nexus analysis of CSL binding to DNA. The results showed a global decrease in Cbf12DBM-TAP binding to its known target loci, however, the ability of Cbf12DBM-TAP to bind DNA was not fully abolished *in vivo* (**Fig. S3A**). This is in contrast to the previous *in-vitro* studies of plasmid-expressed HA-Cbf12, where binding to DNA was no longer detectable upon the introduction of the DBM mutation (Oravcová et al., 2013). We visually inspected ChIP-nexus read coverage at the known adhesion-related genes and identified *cbf12* and flocculins *gsf2*, *pfl7* and *pfl2* as potential targets of Cbf12 (**Fig. 1D**, **Fig. S3B**). Interestingly, with the exception of *pfl2*, these genes were previously found to be upregulated upon Cbf12 overexpression (Kwon et al., 2012). Nevertheless, of these four loci, only *pfl7* and *pfl2* showed markedly decreased CSL binding in the *cbf12DBM-TAP* mutant (**Fig. 1D**, **Fig. S3B**). Subsequent ChIP-qPCR and RT-qPCR analyses aimed at the *cbf12*, *gsf2* and *pfl7* genes confirmed the ChIP-nexus results (**Fig. 1E**) and showed that only the expression of *pfl7*, but not of *cbf12* and *gsf2*, was decreased in the *cbf12DBM-TAP* mutant (**Fig. 1C**). A similar expression pattern was observed in the *Δcbf12* strain (**Fig. 1C**). Thus, while our results clearly support *pfl7* (and possibly *pfl2*) as a direct target of Cbf12 regulation, the relationship between Cbf12 and *gsf2* and the mechanism of *cbf12* autoregulation remain unclear. Likewise, it still needs to be determined how the antagonism in the regulation of adhesion between Cbf11 and Cbf12 is achieved, even though Cbf11 likely acts by affecting Cbf12. In any case, the ability to bind DNA is critical for CSL proteins to regulate cell adhesion.

### Cbf11DBM cells recapitulate the mitotic and cellular morphology defects of *Δcbf11* cells

Cells lacking Cbf11 display defects in cell morphology (misshapen, enlarged, branched cells), septation defects (multiple septa, thick septa, filamentous growth), and are prone to catastrophic mitosis (“cut” phenotype) (Prevorovský et al., 2009) (**Fig. 2**). To assess the impact of the DBM mutation on the role of Cbf11 in cell cycle regulation, we performed microscopy of WT, *cbf11-TAP*, *Δcbf11* and *cbf11DBM-TAP* cells. While the *cbf11-TAP* cells resembled WT cells, the *cbf11DBM-TAP* mutant largely recapitulated the defects of *Δcbf11* cells. Namely, the *cbf11DBM-TAP* strain showed enlarged and misshapen cells, short multicellular filaments, decondensed and/or fragmented nuclei (**Fig. 2A**), and increased incidence of catastrophic mitosis (**Fig. 2A, B**).

**Figure 2:**
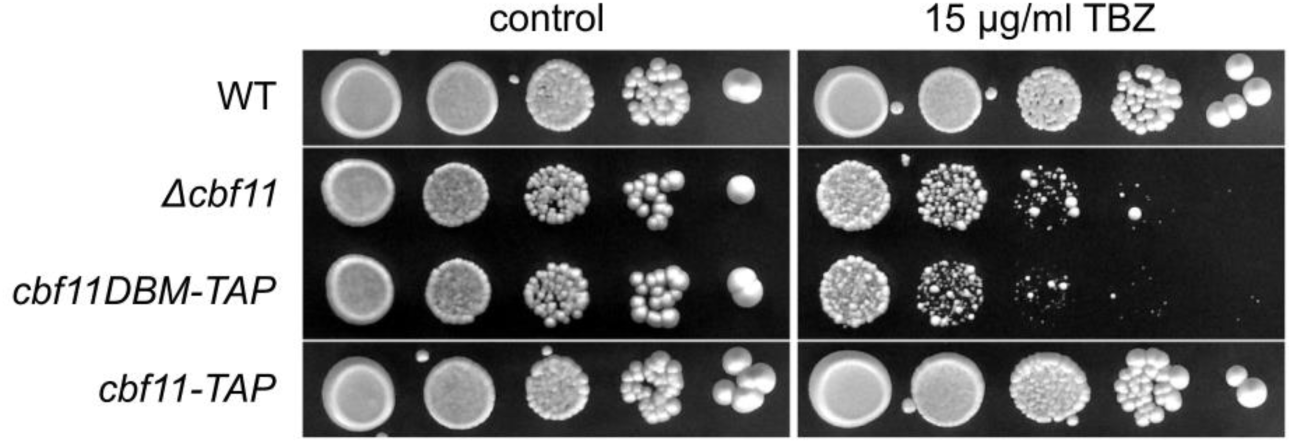
Cbf11DBM cells display the same mitotic and morphology defects as *Δcbf11* cells. **A)** Microscopy of ethanol-fixed, DAPI-stained (nuclei) cells from exponential phase. Representative images from 4 independent experiments. *cbf11DBM-TAP* cells resemble *Δcbf11*: aberrant morphology of cells and nuclei, fragmentation of the nuclear mass (cyan asterisks), “cut” phenotype (yellow asterisks). Scale bar represents 10 μm. **B)** Quantification of the occurrence of the “cut” phenotype in cells from panel A. Both *Δcbf11* and *cbf11DBM-TAP* mutants display the “cut” phenotype with similar penetrance. Mean ± SD values, as well as individual data points for ≥4 independent experiments are shown. Significance was determined by a two-tailed unpaired t-test.

The *Δcbf11* mutant had been found to be sensitive to the microtubule poison thiabendazole (TBZ) in a high-throughput screen (Han et al., 2010). We recently confirmed this observation, and detected aberrant chromatin structure and cohesin occupancy at the centromeres in *Δcbf11* cells (Vishwanatha et al., 2023). When we now tested the *cbf11DBM-TAP* strain, it showed TBZ sensitivity comparable to the *Δcbf11* mutant (**Fig. S4**). So in summary, the DNA binding activity is critical for Cbf11 to exert its role in the regulation of cell cycle progression, mitosis and nuclear integrity.

### Intact DNA-binding domain is necessary for Cbf11 to regulate lipid metabolism

Cbf11 activates the expression of several lipid metabolism genes and is required for the formation of storage lipid droplets (LDs) (Převorovský et al., 2015; Převorovský et al., 2016). We therefore tested the requirement for DNA binding of Cbf11 to exert its regulatory function in lipid metabolism. Indeed, introduction of the DBM mutation into the *cbf11* gene resulted in dysregulation of lipid metabolism very similar to that of the *Δcbf11* strain. Namely, the *cbf11DBM-TAP* culture showed pronounced cell-to-cell heterogeneity in the content of lipid droplets (**Fig. 3A**) and, on average, cells contained fewer and smaller LDs compared to WT (**Fig. 3B**). Next, we determined the expression levels of putative Cbf11 target genes involved in lipid metabolism (*cut6*, *ole1*, *lcf1*, *lcf2*, *fsh2*, *vht1*, *bio2*, *fas2*) (Převorovský et al., 2015) by RT-qPCR. With the exception of *fas2* (alpha subunit of the fatty acid synthase complex), all tested genes were significantly downregulated in *cbf11DBM-TAP* cells to a level similar to the *Δcbf11* mutant (**Fig. 3C**).

**Figure 3:**
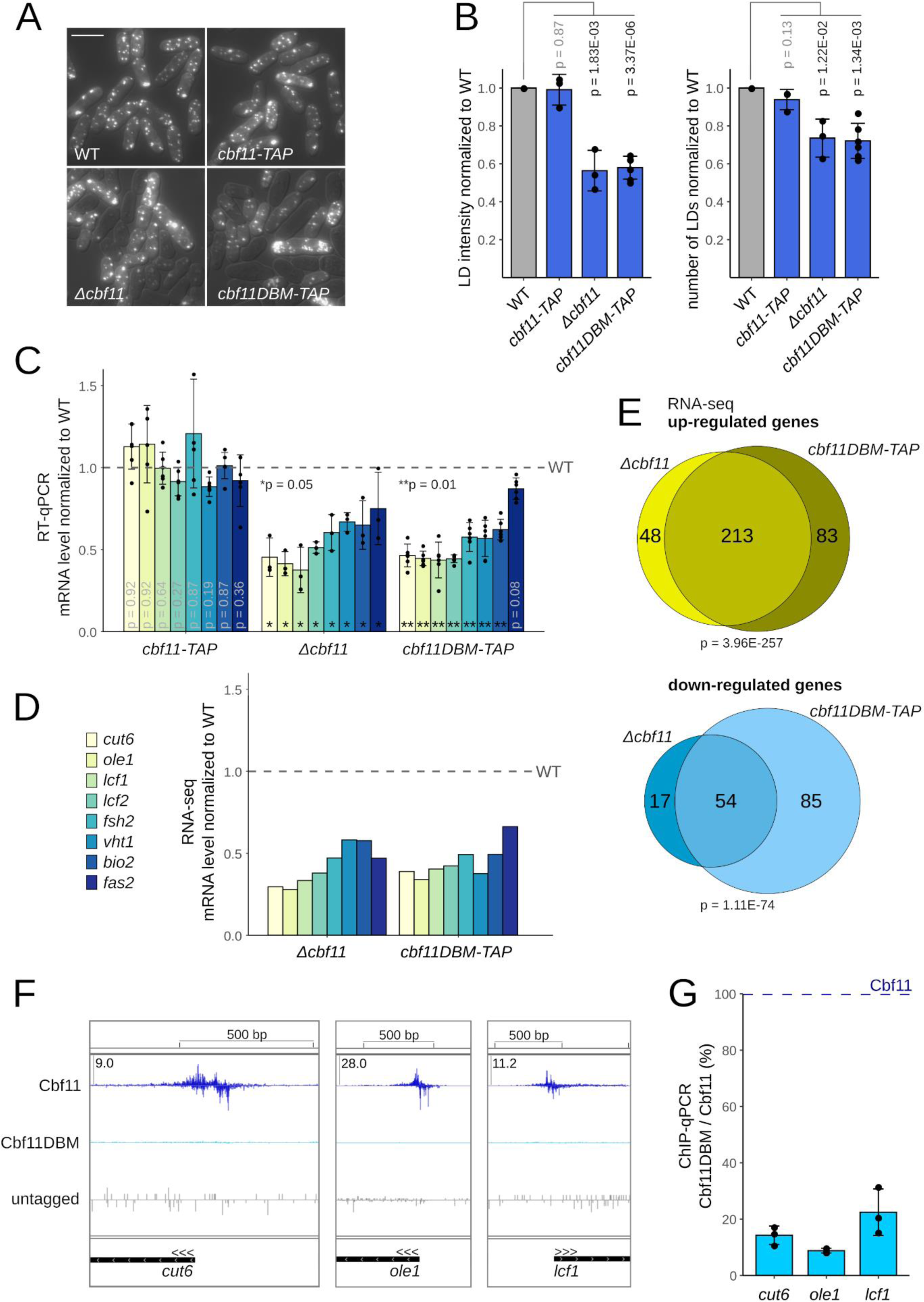
Intact DNA-binding domain of Cbf11 is necessary for its role in lipid metabolism regulation. **A)** Lipid droplet (LD) formation in exponentially growing cells. Both *Δcbf11* and *cbf11DBM-TAP* mutants display decreased LD content and heterogeneity. Representative microscopic images for 3 independent experiments of live cells with LDs stained with BODIPY 493/503. DIC overlay is shown to mark cell boundaries. Scale bar represents 10 μm. **B)** Total fluorescence intensity (left) and number of identified LDs (right) per unit of cell volume. Mean ± SD values, as well as individual data points for ≥3 independent experiments are shown. Significance was determined by a two-tailed unpaired t-test. **C)** Expression of selected lipid-related genes in the indicated cells grown to the exponential phase was analyzed by RT-qPCR. mRNA levels are significantly decreased (except for *fas2*) in *Δcbf11* and *cbf11DBM-TAP* cells compared to WT. Mean ± SD values, as well as individual data points for ≥3 independent experiments are shown. One-sided Mann-Whitney U test was used to determine statistical significance. The legend is shared with panel D. **D)** Expression of the lipid-related genes from panel C determined by RNA-seq. Mean values for 3 independent experiments are shown. **E)** Venn diagram of genes showing differential expression in *Δcbf11* and *cbf11DBM-TAP* cells. Only protein-coding genes showing at least 2-fold and statistically significant change in expression compared to WT are included. Overlap significance was determined using Fisher’s exact test. **F)** *In vivo* binding of Cbf11 and Cbf11DBM TAP-tagged proteins to the promoters of selected lipid-related genes in the exponential phase was analyzed by ChIP-nexus. Mean strand-specific coverage profile of 3 independent experiments for Cbf11 and Cbf11DBM, and a strand-specific coverage profile for untagged WT cells (negative control) are shown. Gene orientation is indicated by three arrowheads; all three sample tracks have the same Y-axis scaling, with the maximum Y-axis value for each locus given in the top-left corner. **G)** ChIP-qPCR shows the decrease in DNA binding of the Cbf11DBM-TAP protein compared to its unmutated counterpart at their selected target loci in the exponential phase. Mean ± SD values, and individual data points of 3 independent replicates are shown.

We then extended the analyses and quantified gene expression globally by RNA-seq of WT, *Δcbf11* and *cbf11DBM-TAP* cells. The RNA-seq results were in good agreement with our previous microarray analyses of the *Δcbf11* transcriptome (**Fig. S5**), and confirmed the ∼50% downregulation of lipid metabolism genes in the *cbf11DBM-TAP* strain (**Fig. 3D**, **Fig. S3C**). Furthermore, there was a strong overlap in the groups of genes expressed differentially in either *Δcbf11* or *cbf11DBM-TAP* cells (**Fig. 3E**). Finally, ChIP-nexus and/or ChIP-qPCR analyses revealed that the decrease in gene expression in *cbf11DBM-TAP* cells coincided with loss of Cbf11DBM binding to the promoters of *cut6*, *ole1*, *lcf1* (**Fig. 3F, G**), and *lcf2*, *fsh2* and *ptl1* genes (**Fig. S3D**). Interestingly, judging by the ChIP-nexus footprints, Cbf11 seems to bind only 1 site per promoter (**Fig. 3F**, **Fig. S3D**). Taken together, the regulatory role of Cbf11 in lipid metabolism requires its ability to bind DNA, and our results confirm Cbf11 as a direct activator of gene expression.

### CSL proteins are required for maintenance of genome integrity

The observations of fragmented and/or decondensed nuclei in *Δcbf11* and *cbf11DBM-TAP* cells (**Fig. 2A**), together with a previous report showing a requirement for the mammalian CSL/RBP-Jκ protein in the maintenance of genome integrity (Bottoni et al., 2019), prompted us to test the sensitivity of CSL mutants to genotoxic stress. Indeed, when exposed to camptothecin (CPT), a topoisomerase I inhibitor causing double-stranded DNA breaks, the *Δcbf11* cells showed clear sensitivity to the drug (**Fig. 4A**). Surprisingly, the *Δcbf12* strain was also sensitive to CPT, albeit to a lesser extent (**Fig. 4A**). To our knowledge, this is the first case where the effects of manipulating the respective CSL paralogs are not antagonistic. Notably, the *Δcbf11 Δcbf12* double mutant showed CPT sensitivity comparable to the *Δcbf11* strain (**Fig. 4A**). The absence of any clear additive effects on CPT sensitivity in the double mutant suggests that both CSL paralogs affect the same process or pathway. Thus, our genetic analyses indicate that Cbf11 is more important for the maintenance of genome integrity than Cbf12, and that Cbf11 likely acts downstream of Cbf12.

**Figure 4:**
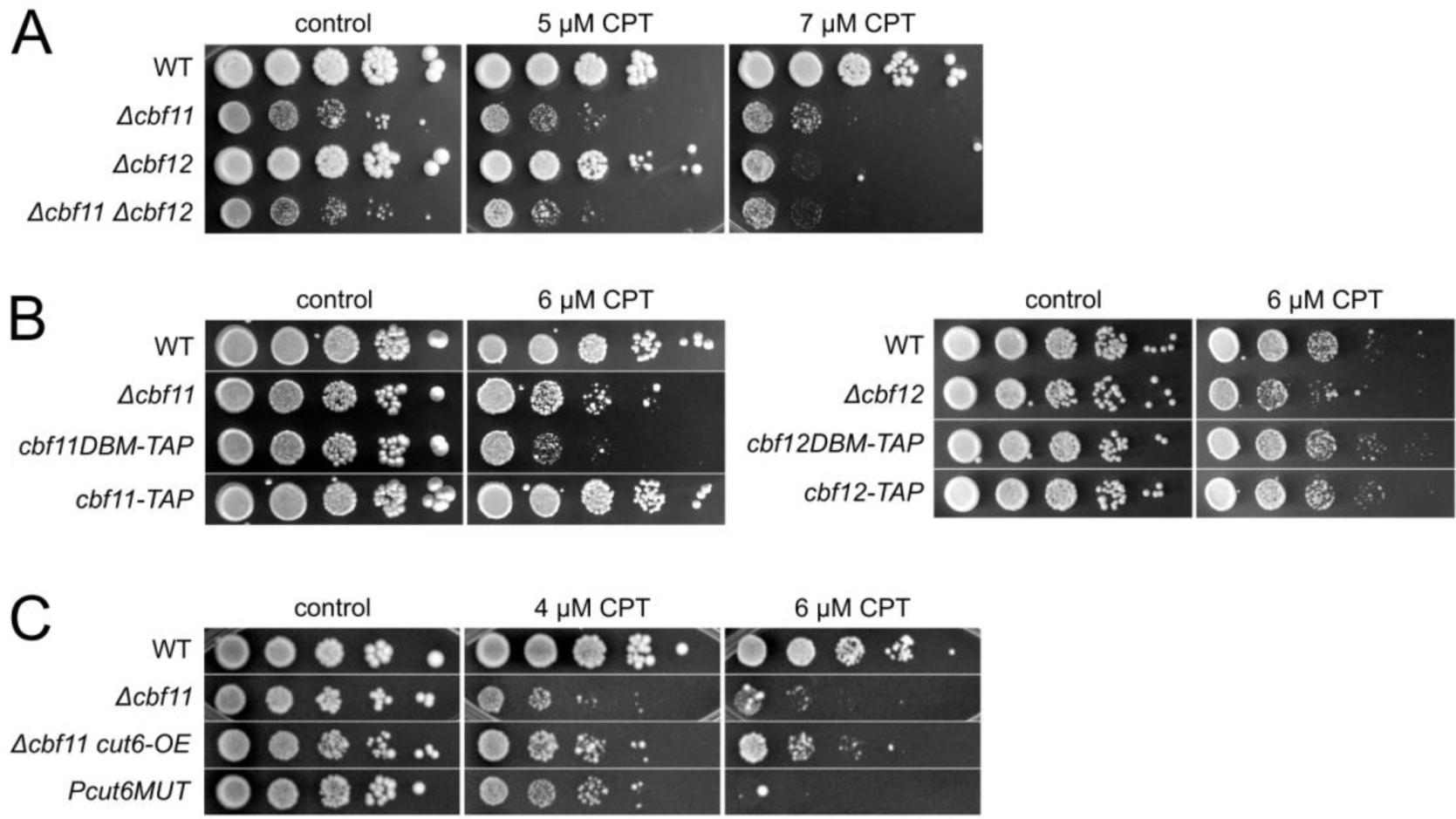
CSL proteins are involved in the maintenance of genome integrity. Cells were spotted in 10-fold serial dilutions on control and testing plates containing indicated amounts of camptothecin (CPT) that causes double-stranded DNA breaks. Cell growth and survival under genotoxic stress was imaged after 5 (panel A), 6 (panel B left), 2 (panel B right) or 4 (panel C) days of incubation. Representative images for 2 independent experiments are shown. **A)** Both *Δcbf11* and *Δcbf12* cells are sensitive to CPT, and the phenotype of the double deletion mutant is not additive. **B)** Cells expressing the Cbf11DBM protein display CPT sensitivity, while cells expressing Cbf12DBM are not sensitive. **C)** *Pcut6MUT* cells are also sensitive to CPT. Mild improvement of *Δcbf11* CPT sensitivity was observed upon *cut6* overexpression (*cut6-OE*).

Next, we tested the impact of CSL DBM mutations on the cellular resistance to genotoxic stress. While the *cbf11DBM-TAP* cells showed CPT sensitivity comparable to *Δcbf11*, the *cbf12DBM-TAP* mutant strain was not sensitive to CPT, in stark contrast to the *Δcbf12* cells (**Fig. 4B**). Thus, the DNA binding activity of Cbf12 is not required for its role in the maintenance of genome integrity, and *cbf12DBM-TAP* is a separation-of-function mutation.

We recently showed that mutations in *cbf11* and several lipid metabolism genes (including *cut6*) result in widespread changes in chromatin structure and in altered expression of oxidative stress-response genes (Princová et al., 2023; Vishwanatha et al., 2023). Therefore, we tested whether there is a more general link between lipid metabolism and the cellular resistance to genotoxic stress. Interestingly, the *Pcut6MUT* mutant, which lacks the Cbf11 binding site in the *cut6* promoter and displays ∼50% reduction in *cut6* expression (Převorovský et al., 2016), was also sensitive to CPT (**Fig. 4C**). This result suggests that unperturbed fatty acid synthesis is indeed important for the cell’s ability to resist genotoxic stress. Moreover, when we overexpressed *cut6* in the *Δcbf11* background, compensating the reduced *cut6* expression in cells lacking Cbf11, the *Δcbf11* sensitivity to CPT was partially rescued (**Fig. 4C**). Therefore, the Cbf11-mediated transcriptional regulation of lipid metabolism is required for the maintenance of genome integrity.

### Non-coding RNAs are upregulated in cells lacking intact DNA binding domain of Cbf11

Global perturbations of chromatin structure can result in aberrant expression of various non-coding RNAs (Anderson et al., 2009; Atkinson et al., 2018). Therefore, we used RNA-seq to characterize the non-coding transcriptomes of cells lacking intact Cbf11. As previously reported (Marguerat et al., 2012), on average, the expression of long non-coding RNAs (lncRNAs) is markedly lower compared to coding genes (**Fig. 5A**), and many annotated lncRNAs might be a mere result of transcriptional noise (Atkinson et al., 2018). Interestingly, hundreds of lncRNAs showed significant changes in expression in both *Δcbf11* and *cbf11DBM-TAP* cells compared to WT, with both the number and amplitude of expression changes being larger in the direction of upregulation (**Fig. 5A**; 1568 up- and 656 downregulated in *Δcbf11*; 1011 up- and 363 downregulated in *cbf11DBM-TAP*). Intriguingly, the changes in lncRNA expression correlated well with the degree of histone H3 lysine 9 acetylation (H3K9ac) at the respective lncRNA loci in the *Δcbf11* mutant (**Fig. 5B**). Therefore, we conclude that the DNA binding activity of Cbf11 is also required for proper expression of the non-coding transcriptome.

**Figure 5:**
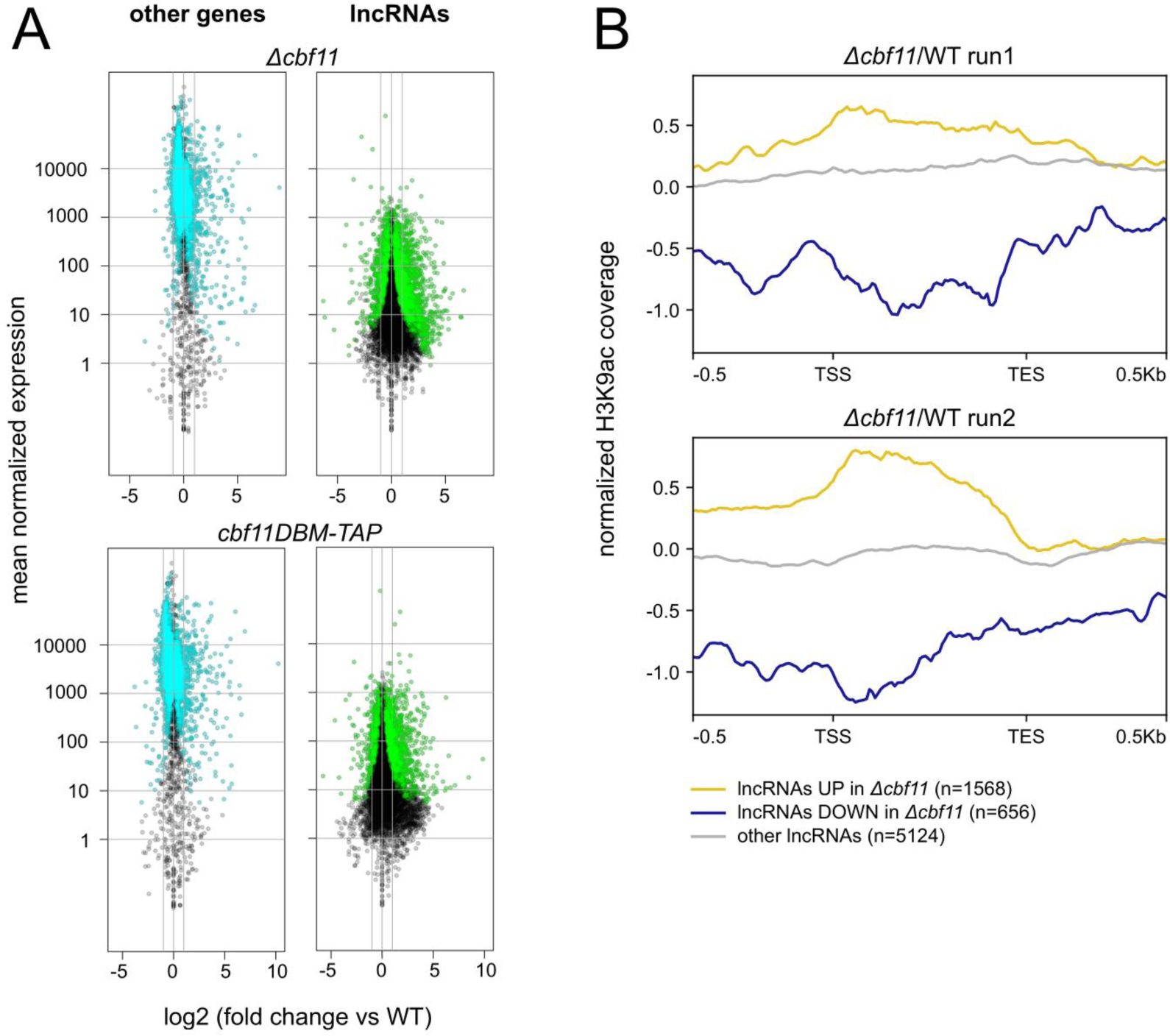
Long non-coding RNAs are upregulated in both *Δcbf11* and *cbf11DBM* cells. **A)** Gene expression in each mutant was determined by RNA-seq and compared to WT. For each gene, the magnitude of change in expression was plotted against its mean expression value across all samples (mutants and WT). Statistically significant differentially expressed genes are shown in cyan (other genes) or green (lncRNAs). The two gray vertical lines indicate the interval of 2-fold up- or downregulation. **B)** Long ncRNA expression in the *Δcbf11* mutant correlates with H3K9 acetylation levels at the corresponding genes. Average-gene profiles of H3K9ac ChIP-seq coverage from 2 independent experiments are shown (Princová et al., 2023). TSS - transcription start site; TES - transcription end site.

## DISCUSSION

The CSL proteins have been implicated in numerous cellular processes in the fission yeast *S. pombe*. Cells lacking CSL genes display a wide range of phenotypes, including altered cell cycle progression, lipid metabolism, resistance to stresses or cell adhesion (Kwon et al., 2012; Prevorovský et al., 2009; Převorovský et al., 2015; Převorovský et al., 2016; Vachon et al., 2013; Vishwanatha et al., 2023). CSL proteins are transcription factors (Oravcová et al., 2013), however, so far only two CSL-regulated processes have been directly linked to their transcription factor activities: activation of lipid metabolism genes by Cbf11 (Převorovský et al., 2016) (**Fig. 3**), and activation of adhesion-related genes by Cbf12 (Kwon et al., 2012) (**Fig. 1**). It was therefore possible that CSL proteins might have additional modes of functioning, independent of their transcription factor activity. Indeed, examples of transcription factors with DNA binding-independent moonlighting roles in other cellular processes have been reported (Lee et al., 2003; Wegrzyn et al., 2009). To shed more light on the ways in which CSL roles are carried out, we investigated the phenotypes of carefully constructed mutants with compromised DNA binding activities of Cbf11 or Cbf12. Our results show that the vast majority of known CSL functions in *S. pombe* depend on their ability to bind DNA (the only exception being the role of Cbf12 in resistance to genotoxic stress). Nevertheless, for many of these roles the exact mechanism(s) of regulation remains to be established.

Notably, most CSL mutant phenotypes are caused by perturbations of the *cbf11* paralog, while *cbf12* seems to be involved almost exclusively in cell adhesion. Importantly, in the present study we show that the DNA binding activity of Cbf11 is implicated in all its known functions, with the regulation of lipid metabolism genes remaining the best characterized role of Cbf11. Strikingly, we showed that perturbed fatty acid synthesis results in large-scale changes in histone modifications (H3K9 acetylation and methylation), which also affect gene expression (Princová et al., 2023; Vishwanatha et al., 2023) (**Fig. 5**). The synthesis of fatty acids consumes acetyl-coenzyme A, a central metabolite that also acts as a substrate for protein acetylation. It is therefore tempting to speculate that changes in chromatin structure resulting from perturbed lipid metabolism and mediated by histone acetyltransferases (Princová et al., 2023) could be the root cause underlying the wide range of defects observed in *cbf11* mutant cells. For example, perturbed centromeric chromatin might be responsible for the altered expression of centromere-proximal genes, decreased mitotic fidelity, and sensitivity to microtubule poisons observed in *Δcbf11* cells (Vishwanatha et al., 2023). In line with this proposition, centromeric heterochromatin was shown to be important for proper centromere function in mitosis (Volpe et al., 2002). Next, mutations in several major fission yeast regulators of chromatin structure are known to compromise genome integrity and lead to sensitivity to genotoxic insults. These regulators include the sole *S. pombe* histone methyltransferase Clr4 and the essential Clr6 histone deacetylase (both target H3K9 and are required for heterochromatin formation), and the HIRA histone chaperone complex (Deshpande et al., 2009; Nicolas et al., 2007; Pan et al., 2012). Finally, perturbed chromatin is generally more prone to aberrant transcription (Anderson et al., 2009; Atkinson et al., 2018), which could explain the striking and widespread upregulation of lncRNA genes in the *cbf11* mutants (**Fig. 5**). Based on these findings, we propose that mere changes in lipid metabolism (such as decreased rate of fatty acid synthesis) may have far reaching and previously unappreciated consequences for the proper functioning of numerous, seemingly unrelated, important cellular processes.

## MATERIALS AND METHODS

### Cultivations, media and strains

The *Schizosaccharomyces pombe* cells were grown according to standard procedures (Petersen and Russell, 2016) in complex yeast extract medium with supplements (YES) at 32°C. Routine optical density (OD) measurements of liquid cell cultures were taken using the WPA CO 8000 Cell Density Meter (Biochrom). Cells were grown to either exponential or stationary phase according to the nature of the experiment. Regarding exponential phase, cells were allowed to reach OD_600_ 0.5. When cultivated to stationary phase, exponentially growing cells were diluted to OD_600_ 0.03-0.15 in fresh medium and cultivated for additional 21-24 hours. Growth rate measurements and calculation of cell culture doubling time (DT) were performed as described previously (Zach et al., 2018). Significance of differences in mutant doubling times was determined by a two-tailed unpaired t-test. Statistical tests were performed on raw data (DT in minutes), however, values normalized to WT were used for plotting to allow for more intelligible visualization. The list of strains used in this study is provided in **Table S1**.

### Construction of strains

Construction of the scarless knock-in strains expressing C-terminally TAP-tagged Cbf11 or Cbf11DBM from its endogenous chromosomal locus was described previously (Princová et al., 2023).

The Cbf12-TAP scarless knock-in strain was constructed by a two-step gene tagging approach using *ura4* selection. First, to replace the 3’ end of *cbf12* ORF in its chromosomal locus, an auxotrophic WT strain (PN559; *ura4-D18*, *leu1-32*, *ade6-M216*, *h^-^*) was transformed using the standard lithium acetate method (Bähler et al., 1998) with a plasmid carrying the *ura4^+^* sequence surrounded by *cbf12* fragments used for homologous recombination (plasmid pMaP21 digested by SalI, KasI, and AlwNI). After selection on EMM+ade+leu plates, the integration of *ura4^+^* into the *cbf12* genomic locus was verified by PCR (primers MaP127+MP31+MP32). Then, the obtained MaP178 strain (*cbf12::ura4^+^*, *ura4-D18*, *leu1-32*, *ade6-M216*, *h^-^*) was used for a second integration step where a TAP-tag was introduced instead of the *ura4^+^* sequence. MaP178 cells were transformed with a fragment of the *cbf12-TAP* sequence (plasmid pMaP29 digested by SalI, KasI, and AlwNI) as template for homologous recombination. Transformants were selected on YES plates containing 5-fluoroorotic acid (5-FOA, 1 g/L) and the integration of the TAP-tagged fragment into the *cbf12* chromosomal locus was verified by PCR (primers MaP127+MaP117+MP32) and sequencing. Expression of the Cbf12-TAP protein in the resulting strain (MaP183; *cbf12-ctap4*, *ura4-D18*, *leu1-32*, *ade6-M216*, *h^-^*) was verified by western blot with an anti-TAP antibody (Thermo Scientific, CAB1001). Prototrophic Cbf12-TAP strains (MP540/MP541) were then prepared by standard crossing and revalidated.

The Cbf12DBM-TAP scarless knock-in strain was constructed and validated analogously with some modifications. To introduce the DNA-binding mutation (DBM; R644H) (Oravcová et al., 2013) into the *cbf12* endogenous locus, MaP178 cells were transformed according to (Okazaki et al., 1990) with a fragment of the *cbf12DBM-TAP* sequence (plasmid pMaP25 digested by SalI and KasI) as template for homologous recombination. Transformants were selected on PMG+ade+leu+ura+5-FOA plates. The integration of the *cbf12DBM-TAP* fragment into the *cbf12* chromosomal locus and the presence of DBM were verified by PCR (primers MaP169+MP32 and chkR+chkF, respectively) and sequencing. Verification of the Cbf12DBM-TAP protein expression in the resulting MaP187 strain (*cbf12DBM-ctap4*, *ura4-D18*, *leu1-32*, *ade6-M216*, *h^-^*) and preparation of the final prototrophic Cbf12DBM-TAP strains (MP518/MP519) were performed as described above for Cbf12-TAP.

The lists of plasmids and primers used in this study are provided in **Table S2** and **Table S3**, respectively.

### Spot tests

Exponentially growing cells were 10-fold serially diluted and spotted onto control YES plates and YES plates containing camptothecin (CPT; 4-7 μM) or thiabendazole (TBZ; 15 μg/ml) for drug sensitivity assays. The spots were allowed to dry, plates were incubated at 32°C and imaged at the day specified in the figure legend.

### Washing assays

Washing assays were used to study cell adhesion. Exponentially growing cells were spotted on an agar plate, incubated at 32°C for 7-8 days, and then washed carefully and evenly with a stream of water. Whole cell spots were imaged before and after washing in **Fig. 1A**. As for **Fig. 1B**, microscopic pictures of the edge of one colony were taken before and after washing.

### Fluorescence microscopy

For observation of nuclear morphology and quantification of the occurrence of the “cut” phenotype, exponentially growing cells were fixed in 70% ethanol, rehydrated in water, stained with 0.1-1 μg/mL DAPI and photographed using the Olympus CellR system or the Leica DM750 microscope. Frequency of “cut” phenotype occurrence was determined by manual counting of “cut” cells using the ImageJ software, version 1.53c (Schneider et al., 2012). 200-1000 cells per sample were analyzed.

Analysis of lipid droplets (LDs) content and automated image processing was performed as described previously (Princová et al., 2019). Briefly, exponentially growing live cells were stained with BODIPY 493/503 (0.1 mg/mL; Invitrogen) and Cascade Blue dextran (10 mg/mL; Invitrogen) to visualize LDs and determine cell boundaries, respectively. Cells were subjected to z-stack imaging using the Olympus CellR microscope. 10-16 images per sample were then processed by an automated pipeline implemented in the MATLAB software. Resulting output data with detected cell objects and LDs were processed using R. A two-tailed unpaired t-test was used to determine statistical significance. Statistical tests were performed on raw data (dotIntensity_per_cellVolume or dots_per_cellVolume, respectively), however, values normalized to WT were used for plotting to allow for more intelligible visualization. Representative microscopic pictures of LDs were prepared by transforming z-stacked images into maximum projection and applying DIC overlay to show cell boundaries in ImageJ ver. 1.53c (Schneider et al., 2012).

### RT-qPCR

Total RNA was extracted from cells at exponential (**Fig. 3C**) or stationary (**Fig. 1C**) phase using the MasterPure Yeast RNA Purification Kit including a DNase treatment step (Biosearch Technologies), and converted to cDNA using random primers and the RevertAid Reverse Transcriptase kit (Thermo Fisher Scientific) following manufacturer’s instructions. Quantitative PCR was performed in technical triplicates using the 5x HOT FIREPol EvaGreen qPCR Supermix (Solis Biodyne) and the LightCycler 480 II instrument (Roche). For RT-qPCR normalization, *act1* (actin) and *rho1* (Rho1 GTPase) were used as internal reference genes. One (**Fig. 3C**) or two-sided (**Fig. 1C**) Mann-Whitney U test was used to determine statistical significance. All statistical tests were performed on data normalized only to internal reference genes (not to WT expression values), however, gene expression values normalized to WT were used for plotting to allow for more intelligible visualization. At least two independent biological experiments were performed. The primers used are listed in **Table S3**.

### ChIP-qPCR

Residual DNA binding of CSL proteins bearing the DBM mutation was measured by ChIP-qPCR. For Cbf11 ChIP-qPCR (**Fig. 3G**), 50 mL cultures of Cbf11-TAP/Cbf11DBM-TAP strains were grown to exponential phase and subsequently fixed with 1% formaldehyde for 30 min and quenched with 125 mM glycine. Regarding Cbf12 ChIP-qPCR (**Fig. 1E**), 3 mL of Cbf12/Cbf12DBM-TAP cultures at stationary phase were brought up to 50 mL with YES and immediately fixed as described above. Cells were washed with distilled water and broken with glass beads in Lysis Buffer (50 mM Hepes pH 7.6, 1 mM EDTA pH 8.0, 150 mM NaCl, 1% Triton X-100, 0.1% sodium deoxycholate, FY protease inhibitors [Serva]) using the FastPrep24 machine (MP Biomedicals). Extracted chromatin was sheared with the Bioruptor sonicator (Diagenode) using 20 cycles of 30 s on, 30 s off at high power settings. An aliquot of chromatin extract was kept as input control, and 0.5-1 mg of chromatin extract was used for overnight immunoprecipitation with 30 μl of BSA-blocked magnetic IgG-coated beads (Invitrogen, 110.41). The precipitated material was washed twice with Lysis Buffer (see above), Lysis 500 Buffer (50 mM Hepes pH 7.6, 1 mM EDTA pH 8.0, 500 mM NaCl, 1% Triton X-100, 0.1% sodium deoxycholate), LiCl/NP-40 Buffer (10 mM Tris-HCl pH 8.0, 1 mM EDTA pH 8.0, 250 mM LiCl, 1% Nonidet P-40, 1% sodium deoxycholate), once in TE (10 mM Tris-HCl pH 8.0, 1 mM EDTA), and eluted in Elution Buffer (50 mM Tris-HCl pH 8.0, 10 mM EDTA, 1% SDS). Cross-links were reversed at 65°C for 6 h, samples were treated with DNase-free RNase A followed by proteinase K digestion, and DNA was purified using standard phenol-chloroform extraction and sodium acetate/ethanol precipitation. Quantitative PCR was performed using the 5x HOT FIREPol EvaGreen qPCR Supermix (Solis Biodyne) and the LightCycler 480 II instrument (Roche) as technical triplicates. Signal (C_t_ values) at *act1* locus (**Fig. 1E**) or geometric average of signals at three loci (*act1*, *sty1*, M40) (**Fig. 3G**) were used as negative controls. Three independent biological repeats were performed. The primers used are listed in **Table S3**.

### RNA-seq

Samples were prepared from 3 biological replicates of *Δcbf11* (MP44) and *cbf11DBM-TAP* (MP712) cells and 4 biological repeats of WT (JB32). Cells were cultured to exponential phase, 10 mL were harvested (600 g, 2 min) and the cell pellet was flash-frozen. Total RNA was isolated using a hot acidic phenol method followed by phenol-chloroform extractions and precipitation (Lyne et al., 2003). Extracted RNA was treated with TURBO DNase (Thermo Fisher Scientific) and purified using Qiagen RNeasy columns. RNA quality was assessed on Agilent Bioanalyzer.

3 samples of each *Δcbf11* and WT were processed at the CCGA (Competence Centre for Genomic Analysis), Christian-Albrechts-University Kiel, Germany. Libraries were prepared using the Illumina TruSeq stranded mRNA Library (poly-A), and sequenced in the pair-end mode on an Illumina NovaSeq 6000 instrument with the NovaSeq 6000 SP Reagent Kit with 100 cycles. The remaining 3 samples of *cbf11DBM-TAP* and 1 sample of WT were processed separately at the Institute of Molecular Genetics (IMG, Czech Academy of Sciences, Prague). The sequencing libraries were synthesized using KAPA mRNA HyperPrep Kit (Illumina platform) (Roche, KK8581) and analyzed on an Illumina NextSeq 500 instrument using the NextSeq 500/550 High Output Kit v2.5 (75 Cycles) (Illumina) with single-end, 75 bp, dual index setup.

### RNA-seq data analysis

The reference *S. pombe* genome and annotation were downloaded from PomBase (2022-05-30; https://www.pombase.org/; (Lock et al., 2019; Wood et al., 2002)). Read quality was checked using FastQC version 0.11.9 (https://www.bioinformatics.babraham.ac.uk/projects/fastqc/). Adapters and low-quality sequences were removed using Trimmomatic 0.39 (Bolger et al., 2014). Clean reads were aligned to the *S. pombe* genome using HISAT2 2.2.1 (Kim et al., 2015) and SAMtools 1.13 (Bonfield et al., 2021; Li et al., 2009). Read coverage tracks were then computed and normalized to the respective mapped library sizes using deepTools 3.5.1 (Ramírez et al., 2016). Mapped reads and coverage data were inspected visually in the IGV 2.9 browser (Robinson et al., 2011). Differentially expressed genes were detected using the RUVseq and DESeq2 packages (Love et al., 2014; Risso et al., 2014) in R/Bioconductor (Gentleman et al., 2004; Huber et al., 2015).

### ChIP-nexus

For Cbf11-related analyses, 400 mL cultures of Cbf11-TAP (MP707), Cbf11DBM-TAP (MP712), untagged WT (JB32, a negative control) and Fkh2-TAP (MP931, forkhead transcription factor unrelated to CSL; a negative control) cells were grown in YES to exponential phase. For Cbf12-related analyses, 10-15 mL of Cbf12-TAP (MP540), Cbf12DBM-TAP (MP518), untagged WT (JB32) and Fkh2-TAP (MP931) cells were grown in YES to stationary phase and were brought up to 400 mL with YES before fixation. Three independent biological experiments were performed for the CSL proteins; the negative controls were grown once for each cultivation condition.

*Saccharomyces cerevisiae* cells expressing the histone H2A.2-TAP (Euroscarf yeast strain collection, original ID SC0421, lab ID MP792) were used as a potential spike-in control for normalization of ChIP samples of TAP-tagged *S. pombe* proteins. *S. cerevisiae* cells were cultivated to exponential phase (OD_600_ 1.8) in the complex YPAD medium at 30°C, and chromatin extract preparation was performed as described below for *S. pombe* with these exceptions: fixation for 30 min, 15 cycles of chromatin shearing. However, after assessing the performance of the spike-in control, we decided not to take it into account during subsequent data analyses.

Cells were fixed by adding formaldehyde to the final concentration of 1%. After 10 min incubation, the remaining formaldehyde was quenched by 125 mM glycine for 10 min. Cells were washed with PBS and broken with glass beads in the Lysis Buffer (50 mM Hepes pH 7.6, 1 mM EDTA pH 8.0, 150 mM NaCl, 1% Triton X-100, 0.1% sodium deoxycholate, FY protease inhibitors [Serva], 1 mM PMSF) using the FastPrep24 machine (MP Biomedicals). Extracted chromatin was sheared with the Bioruptor sonicator (Diagenode) using 20 cycles of 30 s on, 30 s off at high power settings. An aliquot of each chromatin extract was kept as the input control sample (used for a qPCR validation step), and 1.5-2 mg of chromatin extract with added *S. cerevisiae* spike-in chromatin (40,000-fold dilution) were used for immunoprecipitation with 100 μl of BSA-blocked magnetic IgG-coated beads (Invitrogen, 110.41). Chromatin immunoprecipitation was performed on a rotating wheel overnight at 4°C. The precipitated material was ChIP-exo treated according to the updated, simplified version of the ChIP-nexus approach (ChIP-nexus [version 2019]; https://research.stowers.org/zeitlingerlab/protocols.html) introduced by (He et al., 2015), except for wash buffers A-D that were optimized for yeasts as previously described (Doris et al., 2018). ChIP-exo treated material was eluted in Elution Buffer (50 mM Tris-HCl pH 8.0, 10 mM EDTA, 1% SDS). Cross-links were reversed at 65°C for 6 h, samples were treated with DNase-free RNase A followed by proteinase K digestion, and purified using standard phenol-chloroform extraction and sodium acetate/ethanol precipitation. qPCR was performed to check the efficiency of ChIP. ChIP-nexus library construction for Illumina sequencing was performed as described in the updated version of the ChIP-nexus approach (see above). Concentration of ChIP-nexus libraries was measured using the Quantus fluorometer (Promega) and fragment size distribution was checked on Agilent Bioanalyzer using the High Sensitivity DNA Assay. Libraries were pooled and sequenced in two technical-repeat runs on an Illumina NextSeq 500 instrument (75 nt single-end mode) using the NextSeq 500/550 High Output Reagent Cartridge v2 (75 cycles) by the Genomics and Bioinformatics service facility at the Institute of Molecular Genetics (Czech Academy of Sciences, Prague). The list of used ChIP-nexus oligonucleotides is provided in **Table S4**.

### ChIP-nexus data analysis

The reference *S. pombe* and *S. cerevisiae* genome and annotation were downloaded from PomBase (2022-05-30; https://www.pombase.org/; (Lock et al., 2019; Wood et al., 2002)). Read quality was checked using FastQC version 0.11.9 (https://www.bioinformatics.babraham.ac.uk/projects/fastqc/). Adapters and low-quality sequences were removed using Trimmomatic 0.39 (Bolger et al., 2014). Reads without an intact, correctly positioned nexus barcode were discarded. Clean reads were aligned to a hybrid *S. pombe & S. cerevisiae* genome using HISAT2 2.2.1 (Kim et al., 2015) and SAMtools 1.13 (Bonfield et al., 2021; Li et al., 2009). PCR duplicates were removed based on unique molecule identifiers (UMI) and mapping information. Read coverage tracks were then computed and normalized to the respective mapped library sizes using deepTools 3.5.1 (Ramírez et al., 2016). Mapped reads and coverage data were inspected visually in the IGV 2.9 browser (Robinson et al., 2011). Peak calling was performed using MACS3 (version 3.0.0a7) (Zhang et al., 2008). Only reproducible peaks present in all three replicates, but absent from the negative control ChIPs (untagged WT, Fkh2-TAP) were kept.

## DATA AVAILABILITY

Source data for Figures, including the numbers of independent experiments (n) in bar plots (**Fig. 1C**, **2B**, **3B**, **3C**, **Fig. S1**) are available as **Supplementary File 2**.

The sequencing data are available from the ArrayExpress database (https://www.ebi.ac.uk/arrayexpress/) under the accession numbers E-MTAB-13305 and E-MTAB-13302 (RNA-seq), and under the accession number E-MTAB-13258 (ChIP-nexus), respectively. Previously published ChIP-seq data (H3K9 acetylation in lipid metabolism mutants) (Princová et al., 2023; Vishwanatha et al., 2023) were obtained from ArrayExpress (accession numbers E-MTAB-11081, E-MTAB-11082).

The scripts used for sequencing data processing and analyses are available from https://github.com/mprevorovsky/CSL-DBM-paper.

## ACKNOWLEDGEMENTS

We would like to thank the Yeast Genetic Resource Center Japan (NBRP Japan) for providing the *fkh2-TAP* strain (FY32982) and the Euroscarf collection center (distributed by Scientific Research and Development GmbH; Germany) for providing the *S. cerevisiae H2A.2-TAP* strain (SC0421). We are grateful to Štěpánka Hrdá and Blanka Hamplová from the Genomics core facility at Biocev Research Center, Faculty of Science, Charles University for Sanger sequencing and quality control of nucleic acids; Valeria Buccheri for help with optimizing the ChIP-nexus protocol; Adéla Kracíková, Kateřina Svobodová and Eva Krellerová for excellent technical assistance; all members of the GenoMik and ReGenEx groups for their support and insightful discussions. Microscopy was performed in the Vinicna Microscopy Core Facility co-financed by the Czech-BioImaging large RI project LM2023050. Computational resources were supplied by the project “e-Infrastruktura CZ” (e-INFRA LM2018140) provided within the program Projects of Large Research, Development and Innovations Infrastructures.

---

**Supplementary Figure S1:**
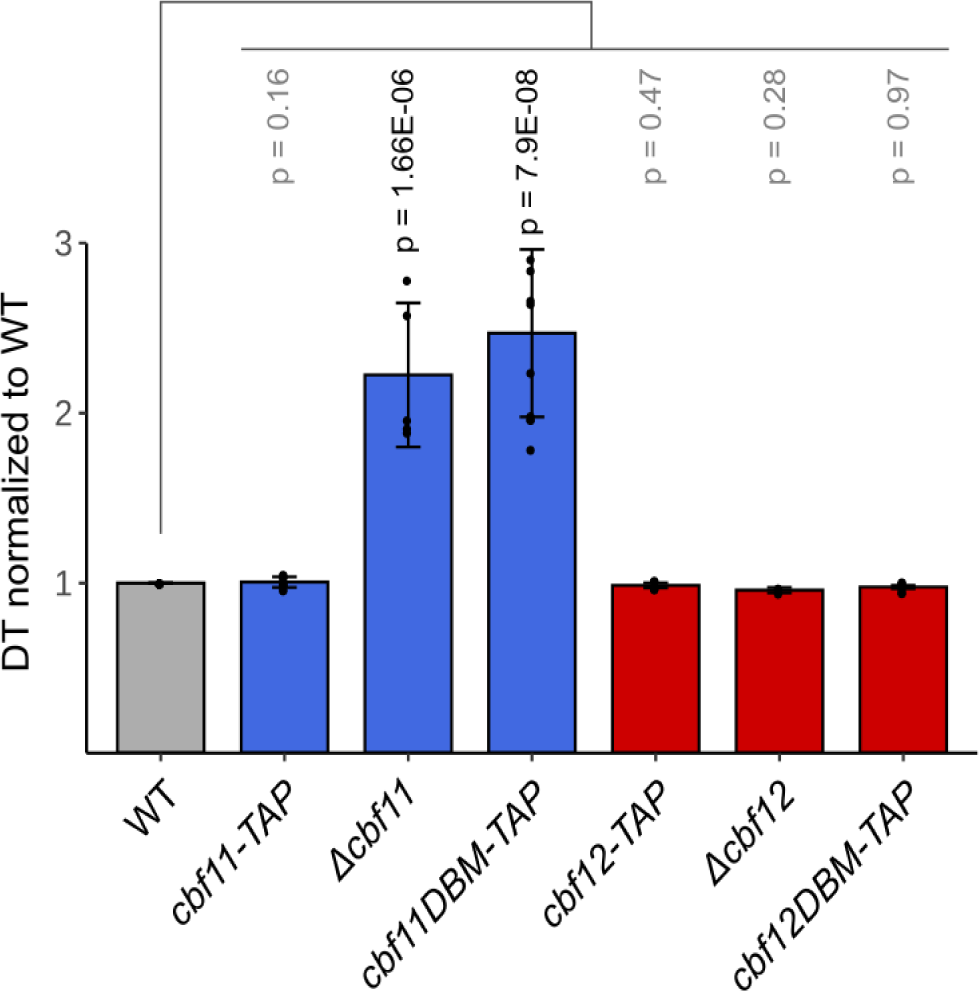
The doubling times of CSL knock-outs and scarless knock-ins. The DBM mutation in Cbf11 increases the time required for biomass doubling in the same manner as *cbf11* gene deletion does. Doubling time (DT) is not affected by any genetic manipulations of the *cbf12* locus. Mean ± SD values, as well as individual data points for ≥3 independent experiments are shown. Significance was tested by a two-tailed unpaired t-test.

**Supplementary Figure S2:**
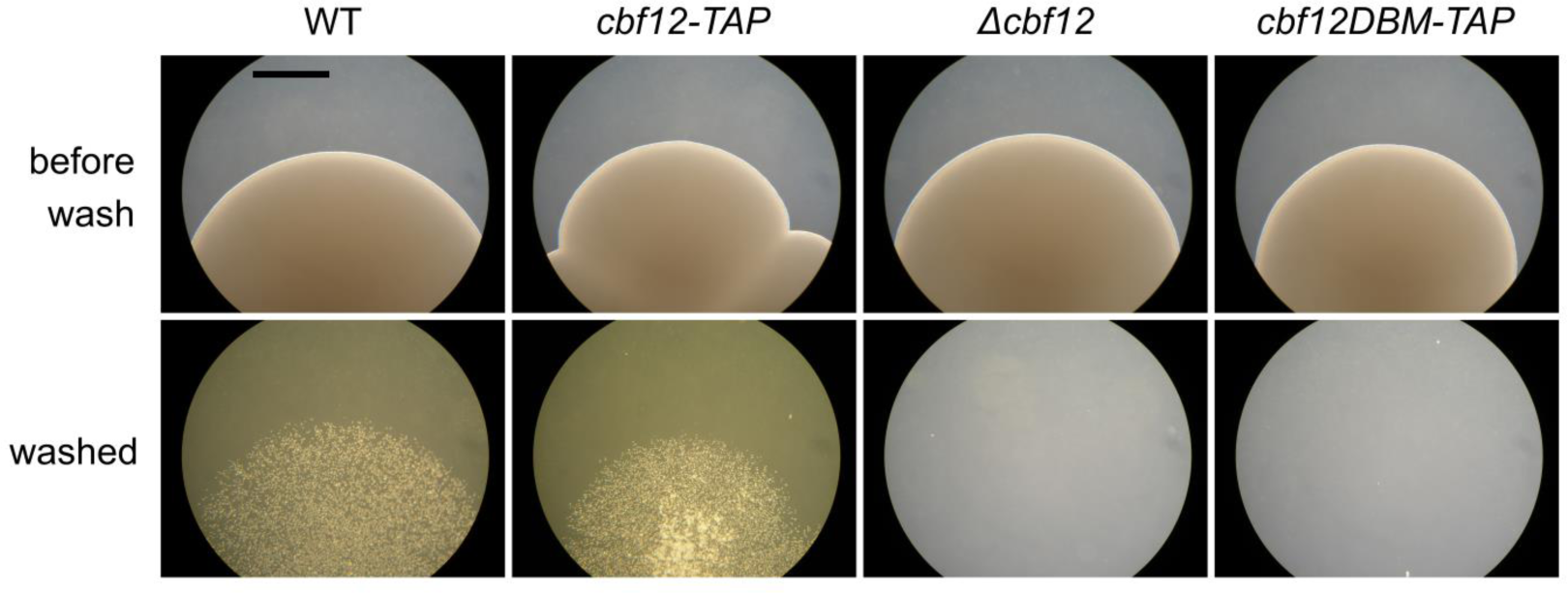
Intact DNA-binding domain of the Cbf12 protein is required for its adhesion-related function. Full non-adjusted microscopic pictures of the washing assay from Fig. 1B are shown. Scale bar represents 0.5 mm.

**Supplementary Figure S3:**
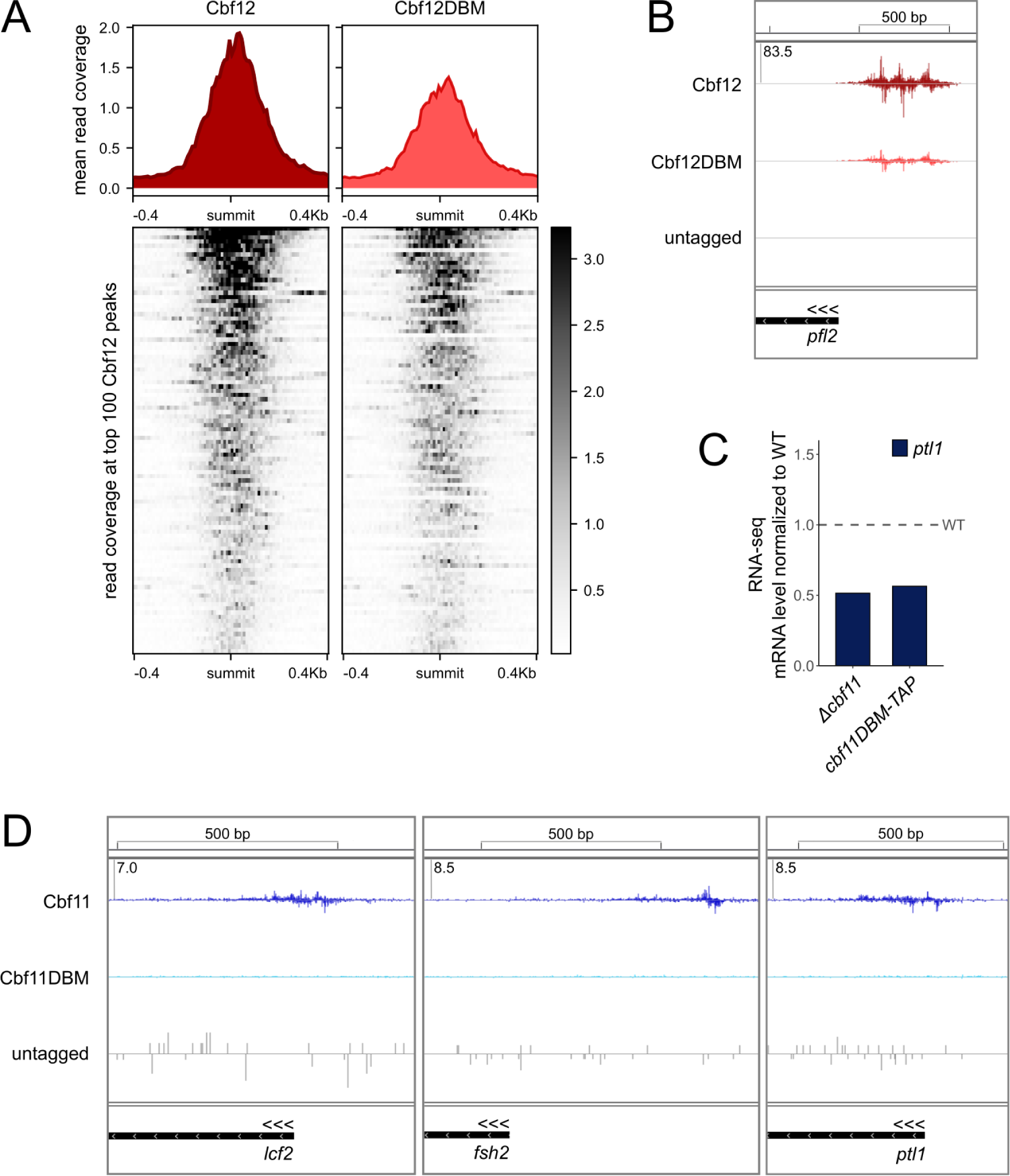
ChIP-nexus and RNA-seq analyses of Cbf11 and Cbf12. **A)** Compared to Cbf12, the binding of the Cbf12DBM protein to DNA is reduced genome-wide. Average-peak profiles of ChIP-nexus coverage (top) and heatmap of coverage at top 100 individual Cbf12 binding sites (bottom) are shown for Cbf12 and Cbf12DBM in the stationary phase. **B)** *In vivo* binding of Cbf12 and Cbf12DBM to the *pfl2* promoter was analyzed by ChIP-nexus. Mean strand-specific coverage profile of 3 independent experiments for Cbf12 and Cbf12DBM, and a strand-specific coverage profile for untagged WT cells (negative control) are shown. Gene orientation is indicated by three arrowheads; all three sample tracks have the same Y-axis scaling, with the maximum Y-axis value given in the top-left corner. **C)** Expression of the lipid-related *ptl1* gene in the indicated cells grown to the exponential phase determined by RNA-seq. Mean values for 3 independent experiments are shown. **D)** *In vivo* binding of Cbf11 and Cbf11DBM to the promoters of additional lipid metabolism genes was analyzed by ChIP-nexus. Mean strand-specific coverage profile of 3 independent experiments for Cbf11 and Cbf11DBM, and a strand-specific coverage profile for untagged WT cells (negative control) are shown. Gene orientation is indicated by three arrowheads; all three sample tracks have the same Y-axis scaling, with the maximum Y-axis value given in the top-left corner.

**Supplementary Figure S4:**
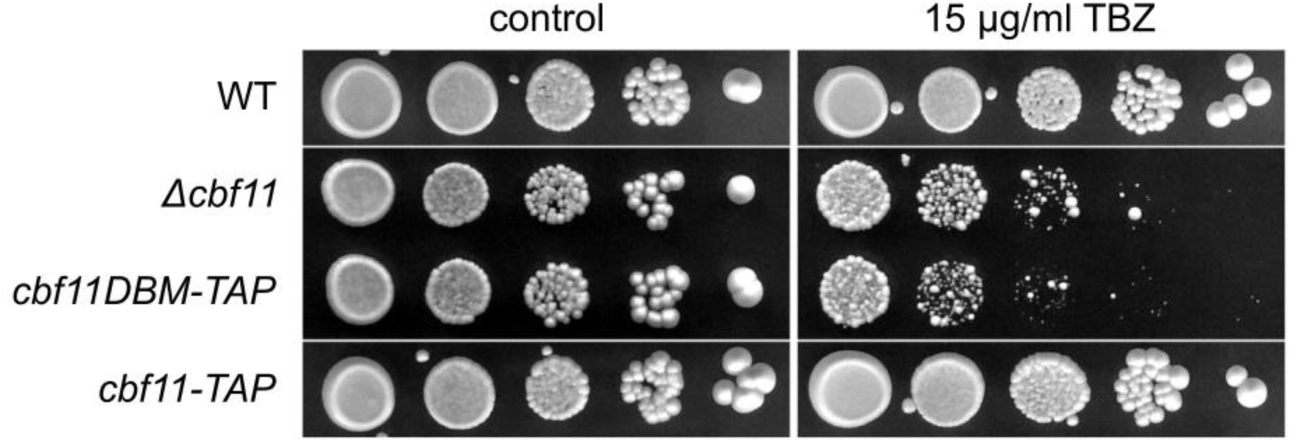
Intact Cbf11 DNA-binding domain is required for resistance to microtubule poison. Both *Δcbf11* and *cbf11DBM-TAP* cells are sensitive to thiabendazole (TBZ), a microtubule-depolymerizing drug. Indicated cultures were spotted in 10-fold serial dilutions on a control plate and plate containing 15 μg/ml TBZ and were grown for 6 days. Representative images of 2 independent experiments are shown.

**Supplementary Figure S5:**
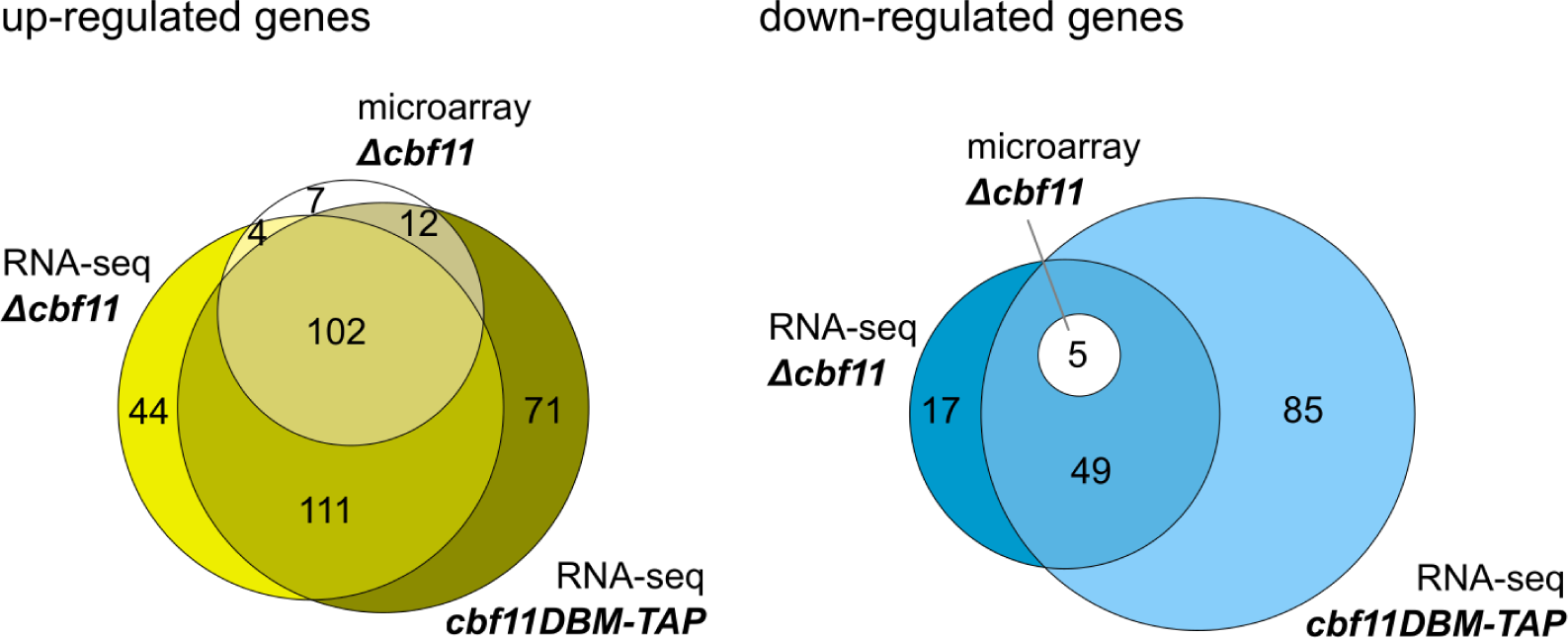
Comparison of Cbf11-related RNA-seq results with previously published expression microarray data. Venn diagram of genes showing differential expression in *Δcbf11* and *cbf11DBM-TAP* as determined by RNA-seq (this study) or microarrays (Převorovský et al., 2015). Only protein-coding genes showing at least 2-fold and statistically significant change in expression compared to WT are included.

